# miniTurbo-based interactomics of two plasma membrane-localized SNARE proteins in *Marchantia polymorpha*

**DOI:** 10.1101/2022.01.21.477208

**Authors:** Katharina Melkonian, Sara Christina Stolze, Anne Harzen, Hirofumi Nakagami

**Affiliations:** Basic Immune System of Plants, Max-Planck Institute for Plant Breeding Research, Carl-von-Linné-Weg 10, 50829 Cologne, Germany; Protein Mass Spectrometry, Max-Planck Institute for Plant Breeding Research, Carl-von-Linné-Weg 10, 50829 Cologne, Germany

**Keywords:** interactomics, Marchantia polymorpha, membrane-trafficking, miniTurbo, proximity labelling, SNARE protein

## Abstract

- *Marchantia polymorpha* is a model liverwort and its overall low genetic redundancy is advantageous for dissecting complex pathways. Proximity-dependent *in vivo* biotin-labelling methods have emerged as powerful interactomics tools in recent years. However, interactomics studies applying proximity labelling are currently limited to angiosperm species in plants.
- Here, we established and evaluated a miniTurbo-based interactomics method in *M. polymorpha* using MpSYP12A and MpSYP13B, two plasma membrane- localized SNARE proteins, as baits.
- We show that our method yields a manifold of potential interactors of MpSYP12A and MpSYP13B compared to a co-immunoprecipitation approach. Our method could capture specific candidates for each SNARE.
- We conclude that a miniTurbo-based method is a feasible tool for interactomics in *M. polymorpha* and potentially applicable to other model bryophytes. Our interactome dataset on MpSYP12A and MpSYP13B will be a useful resource to elucidate the evolution of SNARE functions.

## Introduction

The liverwort *Marchantia polymorpha* is a well- established model plant. The *M. polymorpha* genome has been sequenced (Bowman *et al*., 2017); (Montgomery *et al*., 2020) and genetic tools have been developed (Ishizaki *et al*., 2008; Ishizaki *et al*., 2013; Kubota *et al*., 2013; Ishizaki *et al*., 2015). In *M. polymorpha*, there is no evidence for whole genome duplication during evolution and the number of paralogs for many regulatory genes is rather low in comparison to other model plants (Bowman *et al*., 2017). Accordingly, low genetic redundancy is a useful feature of *M. polymorpha* in dissecting basic mechanisms and gene functions underlying complex pathways.

Elucidating complex pathways and protein-interaction networks remain major challenges in plant research. Conventional approaches to study protein-protein interactions, like co-immunoprecipitation (Co-IP) followed by mass spectrometry (MS) have limitations. Successful enrichment and purification under non-physiological conditions require a certain binding affinity between interactors. Therefore, Co-IP is often effective to capture stable complexes, while weak and transient associations can easily be lost. The analysis of interactions between members of subcellular proteomes may require an enrichment of cellular compartments before Co-IP to avoid artificial interactions upon cell lysis. In this context, proximity-dependent *in vivo* labelling (PL) approaches are gaining an increasing importance as alternative interactomics approaches.

The miniTurbo-based PL method was initially established in bacterial and mammalian cells, and is based on biotin ligase-mediated labelling of interaction partners with exogenously applied biotin (Branon *et al*., 2018). TurboID is a promiscuous biotin ligase that has been engineered from *E. coli* BirA with 15 mutations and has a higher ligase activity at a wide range of temperatures compared to BirA. The miniTurbo is a smaller version of TurboID with a deleted N-terminal domain and has 13 mutations compared to BirA. TurboID and miniTurbo were reported to have higher activities than other biotin-ligases, namely BioID, BioID2, and BASU, in HEK 293T cells. Compared to TurboID, miniTurbo was found to be overall 1.5 times less active and showed a lower background labelling activity without exogenous application of biotin in HEK 293T cells (Branon *et al*., 2018). For PL of interaction partners, the biotin ligase is genetically fused to a bait. Molecules in proximity of the bait are biotinylated by the ligase in the presence of biotin. After labelling, biotinylated proteins can be extracted and enriched by streptavidin-pulldown before an identification by MS.

For pulldown of biotinylated proteins, it is not required that proteins and complexes remain in their native state. Because enrichment of biotinylated proteins does not rely on affinity to the bait proteins. PL approaches can thus also capture weak or transient interactions. The binding affinity between biotin and streptavidin is high. Therefore, protein extraction, binding, and washing steps can be conducted in the presence of high concentrations of detergents. The ligase activity can be inactivated during protein extraction, and thus artificial labelling upon cell lysis will not occur. This is advantageous for investigating interactions of proteins in subcellular compartments, including plasma membrane- localized proteins.

Since biotin ligases do not distinguish between real interaction partners and other molecules residing in proximity of the bait by chance, a certain level of unspecific labelling is expected. Unspecific labelling may occur according to an expression of the bait in a specific cellular compartment, cell- type, tissue, organ, at a specific developmental stage, or physiological status and will be enhanced under saturating labelling conditions. Therefore, non-saturating labelling conditions are desirable and appropriate controls should be designed to narrow down candidates of high confidence by minimizing false positive identifications (Mair *et al*., 2019).

Plants, unlike animals, can synthesise biotin *de novo* (Baldet *et al*., 1993; Baldet *et al*., 1997). High levels of endogenous biotin may lead to background labelling, potentially interfering with PL approaches. The endogenous biotin level in *M. polymorpha* has not been investigated. In *A. thaliana*, the sucrose-H^+^ symporter AtSUC5 was identified to mediate uptake of exogeneous biotin (Ludwig *et al*., 2000; Pommerrenig *et al*., 2013). Until to date, AtSUC5 remains the only sucrose transporter that was shown to function in biotin uptake *in planta*. It is not yet known whether comparable mechanisms for biotin uptake exist in *M. polymorpha*.

The TurboID or miniTurbo method has been successfully applied in *A. thaliana*, *Nicotiana benthamiana*, and *Solanum lycopersicum* using stable transgenic lines or transient expression systems, to efficiently label and identify subcellular proteomes in specific cell types (Mair *et al*., 2019) as well as interacting partners of cytosolic (Zhang *et al*., 2019; Arora *et al*., 2020) and nuclear (Mair *et al*., 2019) bait proteins by MS. Arora *et al*. (2020) demonstrated that TurboID can be applied to detect known interactions of plasma membrane- localized proteins with a targeted approach using immunoblotting. A direct comparison of interactomics applying Co-IP and PL using the same plant materials is still missing. It remains unclear whether biotin ligase-mediated PL approaches can be sensitive and specific to reveal differences in interactomes of very similar proteins, like closely related homologs. Lastly, in bryophyte species, an interactome study utilizing PL approaches has not yet been reported.

In *M. polymorpha*, the two SNARE (soluble N- ethylmaleimide-sensitive factor attachment protein receptor) proteins, MpSYP12A and MpSYP13B, are plasma membrane- localized and ubiquitously expressed throughout the thallus (Kanazawa *et al*., 2016; Kanazawa *et al*., 2020). Plant SNAREs modulate membrane-trafficking, intra- and intercellular signalling, and transport. In *A. thaliana,* 65 SNARE proteins have been identified, 9 of which are SYP1 family proteins that are plasma membrane-localized (Uemura *et al*., 2004). During land plant evolution, the expansion of SNARE proteins and their functional diversification was hypothesized to be linked to multicellularity and likely facilitated the adaptation to a terrestrial lifestyle (Sanderfoot, 2007). Thus, land plant secretory pathways are highly sophisticated, dynamic, and diversely regulated, being involved in a manifold of cellular processes ranging from polarized growth to defence responses (Batoko & Moore, 2001; Collins *et al*., 2003; Catalano *et al*., 2007; Enami *et al*., 2009; Silva *et al*., 2010; Reichardt *et al*., 2011; Uemura *et al*., 2012; Ichikawa *et al*., 2014; Johansson *et al*., 2014; Yun *et al*., 2016; Xia *et al*., 2019; Hirano *et al*., 2020; Rubiato *et al*., 2021). In *M. polymorpha,* the SYP1 protein family is comprised of 4 members: SYP12A, SYP12B, SYP13A, and SYP13B. MpSYP13A and MpSYP13B belong to the SYP13 group and are closely related to AtSYP131 and AtSYP132. MpSYP12A and MpSYP12B belong to the SYP11/12 group, which is phylogenetically separated from the SYP11 or SYP12 group proteins in angiosperms (Kanazawa *et al*., 2016; Bowman *et al*., 2017). Carella *et al*. (2018) reported that MpSYP13B accumulated in haustoria-like structures upon infection of *M. polymorpha* with the oomycete pathogen *Phytophthora palmivora*. Recently, MpSYP12A was shown to localize to the phragmoplast during cell plate formation (Kanazawa *et al*., 2020). Interaction partners of SYP1 family proteins in *M. polymorpha* have not yet been identified.

In this study, we established a miniTurbo-based PL method for interactome profiling in *M. polymorpha*. Using MpSYP12A and MpSYP13B as baits, we evaluated biotin- labelling conditions and a procedure to enrich biotinylated proteins, and then potential interactors were identified by MS. We directly compared the performances of Co-IP and PL approaches using the same plant materials. Lastly, by comparing the identified interactomes of MpSYP12A and MpSYP13B, we found potential interactors that are specific to each SNARE.

## Materials and Methods

### Construction and cloning

Gateway entry vectors containing genomic sequences for an expression of MpSYP12A and MpSYP13B under their own regulatory elements (5′- and 3′- flanking sequences and introns) in *M. polymorpha* were provided by Takashi Ueda (Kanazawa *et al*., 2016). For N- terminal tagging of MpSYP12A and MpSYP13B with miniTurbo and Myc-tag, entry vector backbones were linearized by restriction with SmaI or BamHI enzymes, respectively. Codon-optimized miniTurbo (Fig. S1) was synthesized by Thermo GeneArt and PCR-amplified from a donor plasmid. Gateway entry vectors pMKMM20 (3.5 kb upstream *SYP13B:*5’UTR*:miniTurbo-Myc*-*SYP13B:*3’UTR) and pMKMM21 (3.5 kb upstream *SYP12A:*5’UTR*:miniTurbo- Myc*-*SYP12A:*3’UTR) were generated by in-fusion cloning (HD enzyme mix; Takara Bio). Gateway binary vectors pMKMM22 and pMKMM23 for the expression of miniTurbo-Myc- MpSYP13B and miniTurbo-Myc-MpSYP12A were generated by LR-recombination (LR clonase; Invitrogen) of pMKMM20 or pMKMM21 with pMpGWB301 (Fig. S2).

### *M. polymorpha* lines used in this study

The male *M. polymorpha* Tak-1 ecotype was used as a wildtype. Transgenic lines in Tak-1 background were generated using the cut thallus method (Kubota *et al*., 2013) and *Agrobacterium* strain GV3101 carrying pMKMM22 or pMKMM23. Transformants were selected using Chlorsulfuron and Cefotaxime antibiotics for two generations. Selected transformants were screened for an expression of miniTurbo-Myc-MpSYP12A and miniTurbo- Myc-MpSYP13B fusion-proteins by immunoblotting. Transgenic lines were chosen that displayed similar expression levels of miniTurbo-Myc-MpSYP12A and miniTurbo-Myc- MpSYP13B for further analyses (Fig. S3a). *M. polymorpha* plants were grown and maintained on Gamborg’s B5 (Duchefa; G0209) half strength solid medium containing 1 % plant agar in a walk-in growth chamber under constant white light (50−60 µmol photons LED m^-2^ s^-1^) at 22−24 °C.

### Sample preparation for PL and Co-IP experiments

*M. polymorpha* Tak-1 and transgenic lines were grown from single gemmae on Gamborg’s B5 half strength solid medium containing 0.8 % plant agar for 10 days under constant white light (50−60 µmol photons LED m^-2^ s^-1^) at 22−24 °C. Ten individual 10-day old thalli were pooled per sample. For Co-IP, untreated thalli were sampled immediately and frozen in liquid nitrogen until further processing. For biotin treatment, thalli were transferred into transparent 6- or 12-well plates (Greiner Bio-One; 657160) and submerged in 0−700 µM biotin (Sigma; B4501) solution in water. The thalli were vacuum-infiltrated with biotin solutions for 5 minutes using a desiccator and the samples were incubated for 0−24 hours at room temperature (RT; 22−25 °C) while shaking. After incubation, the thalli were washed once with ice-cold ultrapure water for 2 minutes to remove excess biotin. For sampling, thalli were transferred onto filter paper (Whatman; 1001-085) and left for 10 seconds to drain off excess liquid. Next, the thalli were pressed onto the filter paper for 5 seconds, immediately transferred to fresh tubes containing 2 stainless-steel beads and snap-frozen in liquid nitrogen. The plant material was ground in liquid nitrogen using a mixing mill (MM400, Retsch) for 5 minutes at 30 Hz. For Co-IP, all following steps were conducted at 4 °C. The finely ground powder was mixed with extraction buffer (50 mM Tris pH 7.5, 150 mM NaCl, 10 % glycerol, 2 mM ethylenediaminetetraacetic acid (EDTA), 5 mM dithiothreitol (DTT), 1 % Triton X-100, 1 % Plant Protease Inhibitor; Sigma P9599) and incubated for 30 minutes for protein extraction. For PL samples, the powder was mixed with 500 µl pre-heated SDT buffer (100 mM Tris pH 7.5, 4 % sodium dodecyl sulfate (SDS), 0.1 M DTT) at 95 °C for 5 minutes and then sonicated for 10 minutes. Cell debris was removed from the extracts by two consecutive centrifugation steps (10,000 g, 10 minutes). The protein concentration in cell extracts for Co-IP and PL was determined using the Pierce 660 nm Assay (Thermo Fisher; 22660). Cell extracts with 500 µg of total protein were used as input for Myc-Trap Co-IP and biotin depletion before affinity-pulldown of biotinylated proteins. For biotin-treated thalli, an aliquot of each sample was taken for immunoblotting and whole proteome analysis. Remaining samples were subjected to biotin depletion before pulldown using streptavidin. Biotin dilution series and time course experiments were each done in two independent experiments (Fig. 1c, d; Fig. S4a, b).

**Fig. 1.**
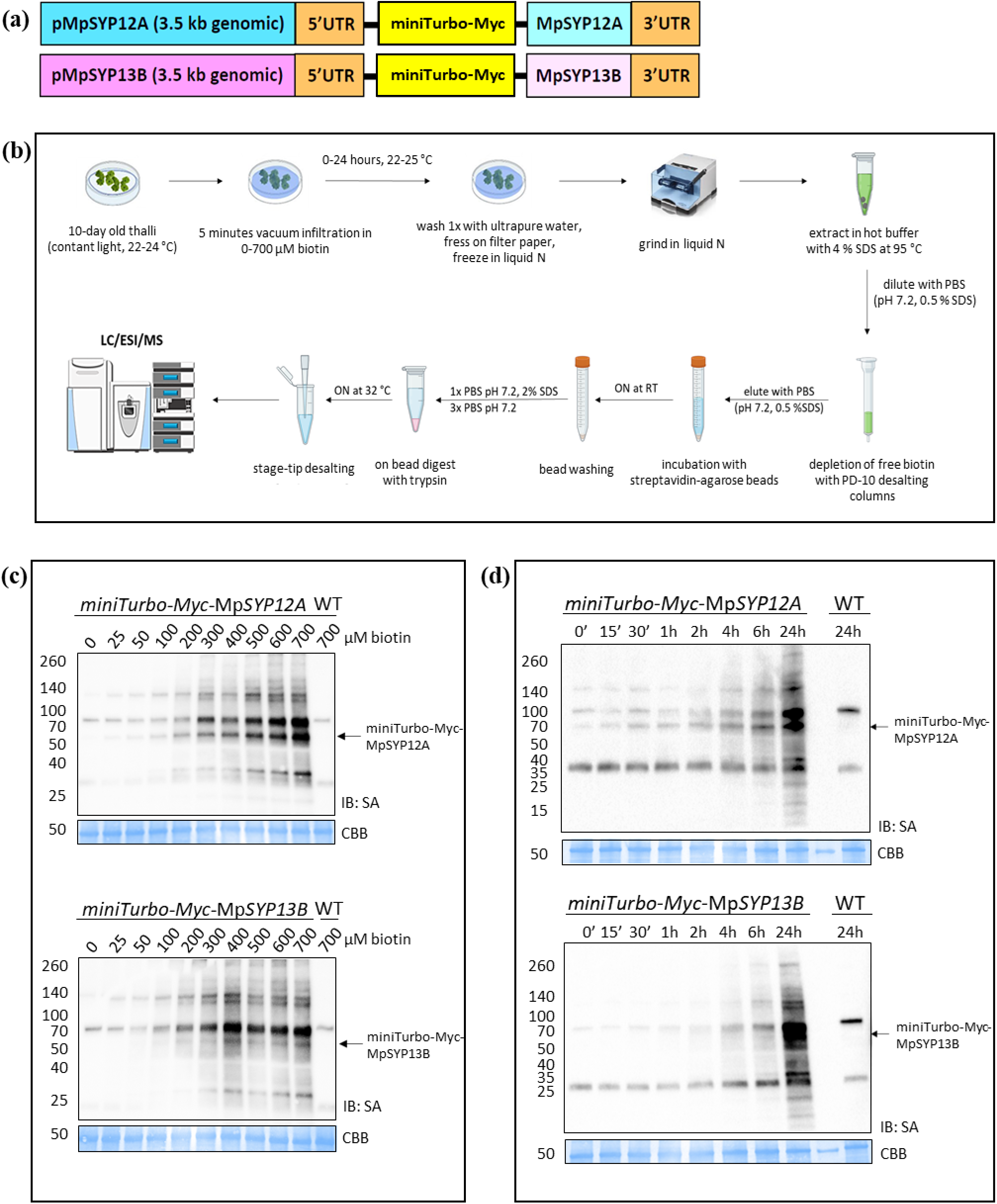
Experimental setup for miniTurbo-mediated biotin-labelling in *M. polymorpha*. (**a**) Schematic representation of the constructs used for generating transgenic plants. (**b**) Overview of the workflow used for evaluating the miniTurbo-based interactomics method in *M. polymorpha*. The figure was created with elements from BioRender (https://biorender.com). (**c** and **d**) Biotin ligase activity in *M. polymorpha*. Streptavidin (SA) immunoblots (IB) of cell extracts from *M. polymorpha* transgenic lines expressing miniTurbo-Myc-MpSYP12A (upper panels) and miniTurbo-Myc-MpSYP13B (lower panels) that were (c) treated with 0−700 µM biotin solutions for 24 hours at RT or (d) treated with 700 µM biotin solution for 0−24 hours at RT. Cell extracts of wildtype Tak-1 (WT) treated with 700 µM biotin solution for 24 hours were used as a control. Arrows indicate the positions of the biotinylated miniTurbo-Myc- MpSYP12A and miniTurbo-Myc-MpSYP13B fusion-proteins. Coomassie Brilliant Blue-stained (CBB) membranes are shown as loading controls.

### Co-IP

Myc-Trap beads (Chromotek; ytma-20) were equilibrated in ice-cold wash buffer (50 mM Tris pH 7.5, 150 mM NaCl, 10 % glycerol, 2 mM EDTA) according to the manufacturer’s instructions. Cell extracts with 500 µg of total protein in 1 ml volume were mixed with 25 µl of equilibrated Myc-Trap beads and pulldown was performed for 2 hours at 4 °C on a rolling wheel. Myc-Trap beads were magnetically separated from the supernatant and washed 3 times in 500 µl wash buffer. An aliquot corresponding to 10 % of the beads was used for immunoblotting and the remaining beads were subjected to on-bead digestion with trypsin. For immunoblotting of Co-IP samples, the proteins were eluted from the beads in 30 µl 4x SDS sample buffer (250 mM Tris-HCl pH 6.8, 40 % glycerol, 8 % SDS, 0.08 % bromophenol blue, 200 mM DTT) by boiling for 10 minutes at 95 °C.

### Depletion of free biotin

Biotin depletion methods were tested using a transgenic line expressing miniTurbo-Myc-MpSYP13B and Tak-1. Cell extracts were prepared as described above from 10-day-old thalli treated with 700 µM biotin for 24 hours. As input, 500 µg of total protein in 500 µl SDT buffer (100 mM Tris pH 7.5, 4 % SDS, 0.1 M DTT) was used per sample. For methanol:chloroform precipitation, 666 µl methanol and 166 µl chloroform were added to the cell extracts and the samples were mixed. Next, 300 µl water was further added and mixed, and then centrifuged for 10 minutes at 4,000 rpm. The separated upper and lower liquid phases were removed. The solid white layer containing precipitated proteins that had formed between the liquid phases was kept. The protein pellet was resuspended in 600 µl methanol and sonicated for 10 minutes. After centrifugation for 10 minutes at 13,000 rpm, the supernatant was removed completely. The protein pellets were air-dried for 5 minutes and resuspended in 500 µl SDT buffer. After 10 minutes sonication, the samples were incubated for 30 minutes at RT while shaking at 1,000 rpm, until the protein pellets were redissolved. The samples were then diluted with PBS buffer (0.1 M phosphate, 0.15 M NaCl, pH 7.2) to a final concentration of 0.5 % SDS. For biotin depletion with PD-10 desalting columns (VWR; 17085101), the columns were equilibrated with SDT:water (1:5) and the cell extracts were diluted to 2.5 ml with ultrapure water. PD-10 desalting was performed according to the manufacturer’s instructions and the proteins were eluted with 3.5 ml PBS buffer containing 0.5 % SDS. For biotin depletion using Zeba spin columns (Thermo; 89893), the columns were equilibrated with SDT:water (1:5) and the cell extracts were diluted to 2.5 ml with ultrapure water. Desalting was then performed according to the manufacturer’s instructions. All biotin-depleted samples were adjusted to 4 ml final volume with binding buffer (0.1 M phosphate, 0.15 M NaCl, 0.5 % SDS, pH 7.2) before pulldown. Aliquots of intermediate steps were taken for immunoblotting. All biotin depletion methods were tested at the same time in duplicates. PD-10 column desalting method was employed for the pulled- down samples measured by MS.

### Pulldown of biotinylated proteins

Biotinylated proteins were pulled-down using streptavidin-agarose beads (Thermo; 20353). Per sample, 100 µl of a 50 % slurry were used. The beads were washed and equilibrated in the binding buffer. Next, the biotin- depleted samples were added to the beads and pulldown was performed overnight at 22−25 °C while mixing. The beads were washed once with 6 ml PBS buffer containing 2 % SDS and then 3 times with 10 ml PBS buffer. Aliquots of all samples were taken during intermediate steps and an aliquot corresponding to 10 % of the washed beads was taken after pulldown for immunoblotting. The proteins were eluted from the beads by boiling for 10 minutes at 95 °C in 50 µl 4x SDS sample buffer containing 20 mM biotin while shaking at 1,000 rpm. For immunoblots, 20 % of the IP-eluate was used, corresponding to 2 % of the input bead-amount. The remaining beads from pulldown with bound biotinylated proteins were subjected to on-bead digestion for MS analysis.

### On-bead digestion

For Myc-IP and streptavidin-pulldown samples, Myc-Trap beads or streptavidin-agarose beads were resuspended in 25 μl digest buffer 1 (50 mM Tris pH 7.5, 2 M urea, 1 mM DTT, 5 µg µl^-1^ trypsin) and incubated at 30 °C for 30 minutes while shaking at 400 rpm. The supernatant was separated from the beads magnetically or by sedimentation and transferred to a fresh tube. The beads were then mixed with 50 µl digest buffer 2 (50 mM Tris pH 7.5, 2 M Urea, 5 mM CAA), and the supernatant was separated from the beads and combined with the supernatant from the previous step. The combined supernatant was further incubated overnight at 32 °C while shaking at 400 rpm. Trypsin was inactivated by acidification with trifluoroacetic acid (TFA), and the peptide sample was subsequently desalted using C_18_ stage tips (Rappsilber *et al*. (2003).

### Sample preparation for whole proteome analyses

Aliquots of cell extracts from biotin-labelling experiments were used for whole proteome analysis. The extracts were processed using a filter-aided sample preparation (FASP) protocol adapted from Wisnewski *et al*. (2009). In brief, 50 µg of total protein extract were used as input. The proteins were alkylated using chloroacetamide and digested using LysC and trypsin. The peptide solutions were desalted using C_18_ stage tips. Whole proteome analyses were conducted for one representative replicate per genotype and condition.

### LC-MS/MS data acquisition

The dried peptides from filter- aided digestion were re-dissolved in buffer A (2 % ACN, 0.1 % TFA) and adjusted to a final peptide concentration of 0.1 µg µl^- 1^ for analysis. The peptide samples from streptavidin- and Myc- Trap pulldowns were dissolved in 10 µl buffer A and measured without dilution.

PL samples were analysed using an EASY-nLC 1200 (Thermo Fisher) coupled to a Q Exactive Plus mass spectrometer (Thermo Fisher). The peptides were separated on 16 cm frit-less silica emitters (New Objective, 75 µm inner diameter), packed in-house with reversed-phase ReproSil-Pur C18 AQ 1.9 µm resin (Dr. Maisch). The peptides were loaded on the column and eluted for 50 minutes using a segmented linear gradient of 5 % to 95 % solvent B (0 minutes: 5 % B; 0−5 minutes -> 5 % B; 5−25 minutes -> 20 % B; 25−35 minutes -> 35 % B; 35−40 minutes -> 95 % B; 40−50 minutes - > 95 % B) (solvent A: 0 % ACN, 0.1 % FA; solvent B: 80 % ACN, 0.1 % FA) at a flow rate of 300 nl per minute. Mass spectra were acquired in data-dependent acquisition mode with a TOP10 method. MS spectra were acquired in the Orbitrap analyser with a mass range of 300−1500 m/z at a resolution of 70,000 FWHM and a target value of 3×10^6^ ions. Precursors were selected with an isolation window of 1.3 m/z. HCD fragmentation was performed at a normalized collision energy of 25. MS/MS spectra were acquired with a target value of 5x10^5^ ions at a resolution of 17,500 FWHM, a maximum injection time of 85 milliseconds and a fixed first mass of m/z 100. Peptides with a charge of 1, greater than 6, or with unassigned charge state were excluded from fragmentation for MS^2^, dynamic exclusion for 20 seconds prevented repeated selection of precursors.

Myc-IP and whole proteome samples were analysed using an EASY-nLC 1000 (Thermo Fisher) coupled to a Q Exactive mass spectrometer (Thermo Fisher). The peptides were separated on 16 cm frit-less silica emitters (New Objective, 75 µm inner diameter), packed in-house with reversed-phase ReproSil-Pur C18 AQ 1.9 µm resin (Dr. Maisch). Peptides (0.5 µg) were loaded on the column and eluted for 115 minutes using a segmented linear gradient of 5 % to 95 % solvent B (0 minutes: 5 % B; 0−5 minutes -> 5 % B; 5−65 minutes -> 20 % B; 65−90 minutes -> 35 % B; 90−100 minutes -> 55 % B; 100−105 minutes -> 95 % B, 105−115 minutes -> 95 % B) (solvent A: 0 % ACN, 0.1 % FA; solvent B: 80 % ACN, 0.1 % FA) at a flow rate of 300 nl per minute. Mass spectra were acquired in data-dependent acquisition mode with a TOP15 method. MS spectra were acquired in the Orbitrap analyser with a mass range of 300–1750 m/z at a resolution of 70,000 FWHM and a target value of 3×10^6^ ions.

Precursors were selected with an isolation window of 2.0 m/z. HCD fragmentation was performed at a normalized collision energy of 25. MS/MS spectra were acquired with a target value of 10^5^ ions at a resolution of 17,500 FWHM, a maximum injection time of 55 milliseconds and a fixed first mass of m/z 100. Peptides with a charge of 1, greater than 6, or with unassigned charge state were excluded from fragmentation for MS^2^, dynamic exclusion for 30 seconds prevented repeated selection of precursors.

### Data analysis

Raw data were processed using MaxQuant software (version 1.6.3.4, http://www.maxquant.org/) (Cox & Mann, 2008) with label-free quantification (LFQ) and intensity based absolute quantification (iBAQ) enabled (Tyanova *et al*., 2016). For PL data, normalization was skipped for the LFQ quantification. MS/MS spectra were searched by the Andromeda search engine against a combined database containing the sequences from *M. polymorpha* (MpTak1v5.1_r1_primary_transcripts_proteinV2; http://marchantia.info/, (Montgomery *et al*., 2020)) and sequences of 248 common contaminant proteins and decoy sequences and the sequence of miniTurbo. Trypsin specificity was required and a maximum of 2 missed cleavages allowed. Minimal peptide length was set to 7 amino acids. Carbamidomethylation of cysteine residues was set as fixed, oxidation of methionine and protein N-terminal acetylation as variable modifications. Peptide-spectrum-matches and proteins were retained if below a false discovery rate (FDR) of 1 %. For PL data, the non-normalized MaxLFQ values of all replicates (4 per condition) were pre-processed in Perseus (version 1.5.8.5, http://www.maxquant.org/) and submitted for normalization analysis using the Normalyzer tool (http://normalyzer.immunoprot.lth.se/, (Chawade *et al*., 2014)). The output was analysed for outliers and 1 replicate per condition was removed. The final data analysis was conducted in MaxQuant as described above on the reduced raw dataset. Statistical analysis of the MaxLFQ values was conducted using Perseus (version 1.5.8.5, http://www.maxquant.org/). Quantified proteins were filtered for reverse hits and hits “identified by site” and MaxLFQ values were log2 transformed.

For PL data, transformed MaxLFQ values were normalized by subtraction of the median per column. After grouping samples by condition only those proteins were retained for subsequent analysis that had 3 or 2 valid values in one of the conditions for PL data or Myc-IP data, respectively. Two-sample *t*-tests were performed using a permutation-based FDR of 5 %. For the generation of volcano plots, missing values were imputed from a normal distribution using the default settings in Perseus (1.8 downshift, separately for each column) for data with 3 valid values in one of the conditions. Volcano plots were generated in Perseus using an FDR of 5 % and an *S0* = 1. The Perseus output was exported and further processed using Microsoft Excel and RStudio (version 1.4.1103, https://www.rstudio.com/), based on R (version x64 4.0.3, https://cran.r-project.org/). The data was processed in RStudio using tidyverse (version 1.3.0), rio (version 0.5.16) and zoo (version 1.8-8) packages. Relative iBAQ values were calculated per column from MaxQuant output, scaled by factor 10^6^ and log10 transformed. The median value of 3 replicates per condition was used to generate volcano plots including relative iBAQ values. Volcano pots were generated in RStudio using the ggplot2 (version 3.3.3), ggrepel (version 0.9.1) and ggsci (version 2.9) packages.

### Immunoblotting

Proteins were separated by SDS- polyacrylamide-gel electrophoresis (PAGE) and blotted onto PVDF membranes (BioRad; 1704272) using a Trans-Blot Turbo (BioRad) transfer system. The following antibodies were used: streptavidin-HRP (Cell Signaling; 3999S), anti-Myc-tag mouse monoclonal antibody (Cell Signaling; 9B11), HRP- conjugated anti-mouse IgG antibody (Cell Signaling; 7076S). The membranes were probed for biotinylated proteins with streptavidin-HRP for 45 minutes to 3 hours at RT. For detection of miniTurbo-Myc, the membranes were probed with anti-Myc-tag primary antibody overnight at 4 °C and then with anti-mouse IgG secondary antibody for 1 hour at RT. Biotinylated proteins or Myc-tagged miniTurbo were visualized on the membranes using a luminol-based chemiluminescent substrate that is oxidized by HRP in the presence of peroxide (Thermo Fisher; 34577). The membranes were stained with Coomassie staining solution (60 mg l^-1^ Coomassie brilliant blue, 10 % acetic acid) afterwards.

### Annotations and gene ontology analyses

For *M. polymorpha* protein annotation, gene annotations from MpTak1_v5.1 and JGI 3.1 (https://marchantia.info/) were integrated. Information of *A. thaliana* homologs was further used for the annotation. A Basic Local Alignment Search Tool (BLAST) was used to determine homologs in *A. thaliana* (TAIR10). The best hit with an e-value ≤ 10^-10^ was defined as a homolog. TAIR10 (https://www.arabidopsis.org/, (Lamesch *et al*., 2012)), PANTHER version 16.0 (http://pantherdb.org/, (Mi *et al*., 2021)), STRING 11.5 (https://string-db.org/, (von Mering *et al*., 2003)), BioGrid 4.4 (https://thebiogrid.org/, (Stark *et al*., 2006)), and IntAct 1.0.2 (https://www.ebi.ac.uk/intact/home, (Orchard *et al*., 2014)) were used for annotating *A. thaliana* homologs (Data S1). RStudio (version 1.4.1103, https://www.rstudio.com/), based on R (version x64 4.0.3, https://cran.r-project.org/), the tidyverse (version 1.3.0), rio (version 0.5.16), and zoo (version 1.8-8) packages, were used for integrating annotation files. GO-term enrichment analysis was performed with Metascape 3.5 (https://metascape.org/, (Zhou *et al*., 2019)) using express analysis settings. Corresponding *A. thaliana* homologs were used as input protein lists for the analysis. A list of reported and predicted interactors of AtSYP1 proteins was generated by integrating information from BioGrid, IntAct, and STRING databases (Data S2). BLAST was used to determine homologs in *M. polymorpha* (JGI 3.1). The best hit with an e-value ≤ 10^-10^ were defined as a homolog (Data S3). PANTHER version 16.0 and PPDB (http://ppdb.tc.cornell.edu/, (Sun *et al*., 2009)) were used to predict plasma membrane-localization of *M. polymorpha* homologs.

## Results

### Experimental design for interactomics using miniTurbo- mediated PL and Co-IP in *M. polymorpha*

To potentially enhance transcription and translation efficiency of the bait-ligase fusion-proteins in *M. polymorpha*, we used a codon-optimized version of the original miniTurbo (Fig. S1). We added a single Myc-tag to the C-terminus of miniTurbo, enabling not only the detection of the miniTurbo fusion- proteins by immunoblotting, but also Co-IP experiments (Fig. 1a, Fig. S1). We designed binary vectors to express miniTurbo- Myc-MpSYP12A and miniTurbo-Myc-MpSYP13B fusion- proteins under the native promoters of Mp*SYP12A* and Mp*SYP13B* genes, respectively (Fig. 1a, Fig. S2). Using these constructs, we generated stable transgenic lines in wildtype Tak-1 background. Expression of the miniTurbo-Myc fusion- proteins in candidate transformant lines was confirmed by immunoblotting using an anti-Myc antibody. To establish and evaluate the miniTurbo-based interactomics method, we selected transgenic lines showing similar expression levels for miniTurbo-Myc-MpSYP12A and miniTurbo-Myc-MpSYP13B. For this study, we used line No. 1 and line No. 3 for miniTurbo-Myc-MpSYP12A and miniTurbo-Myc-MpSYP13B, respectively (Fig. S3a).

In *A. thaliana*, Mair *et al*. (2019) demonstrated that levels of miniTurbo-mediated biotinylation of cellular proteins saturated when plants were treated with 50 µM biotin solution. Since uptake of exogenous biotin and levels of endogenously produced biotin are unknown in *M. polymorpha*, we treated the selected transgenic lines and Tak-1 with different concentrations of biotin, 0−700 µM, to find suitable conditions for *in vivo* biotin-labelling. For biotin treatment, thalli were grown from single gemmae for 10 days, and then whole plants were submerged in biotin solutions and vacuum infiltrated for 5 minutes. We incubated the thalli in biotin solutions at 22−25 °C for 24 hours, a time point at which we expected a saturation of biotin-labelling based on previous studies using miniTurbo in other plant species (Mair *et al*., 2019; Zhang *et al*., 2019). We subsequently checked the levels of biotinylated proteins in cell extracts by immunoblotting using streptavidin-HRP (Fig. 1c, Fig. S4a). We found increasing levels of biotinylated proteins with increasing biotin concentration in cell extracts of transgenic lines expressing either miniTurbo-Myc-MpSYP12A or miniTurbo-Myc-MpSYP13B but not in Tak-1. This confirmed biotin uptake and miniTurbo biotin ligase activity in *M. polymorpha* under the tested conditions. Given that levels of biotinylated proteins did not saturate for biotin concentrations up to 700 µM in *M. polymorpha*, we used 700 µM biotin solution in all following experiments. To determine suitable biotin treatment times in *M. polymorpha*, we next performed a time-course experiment and checked the levels of biotinylated proteins by immunoblotting after 0−24 hours of biotin treatment (Fig. 1d). We detected increased levels of biotinylated proteins in samples of transgenic lines expressing either miniTurbo-Myc-MpSYP12A or miniTurbo-Myc- MpSYP13B after 30 minutes of treatment (Fig. S4b), which further increased over time up to 24 hours of treatment (Fig. 1d).

Previously published studies using TurboID and miniTurbo identified the depletion of free biotin after labelling as a critical step for pulldown of biotinylated proteins using streptavidin beads (Mair *et al*., 2019; Zhang *et al*., 2019; Arora *et al*., 2020; Zhang *et al*., 2021). We therefore tested three different approaches, methanol:chloroform precipitation, PD-10 gravity column desalting, and Zeba spin column desalting, to remove excess free biotin from *M. polymorpha* cell extracts. We used the miniTurbo-Myc-MpSYP13B line, which was treated with 700 µM biotin for 24 hours. We evaluated biotin depletion based on pulldown efficiency, through immunoblotting of biotinylated proteins that could be eluted from the streptavidin-agarose beads after pulldown, and of non-bound biotinylated proteins that remained in the supernatant of the beads (Fig. S3b, c). We found that all three methods could be used before affinity-pulldown to sufficiently enrich biotinylated proteins from *M. polymorpha* samples (Fig. S3b). With respect to easier handling, we used PD-10 column desalting for subsequent experiments.

### The miniTurbo-based approach identifies a manifold of potential interactors of MpSYP12A and MpSYP13B in comparison to the Co-IP approach

A direct comparison between the performances of PL and Co-IP approaches for interactomics using the same plant materials has not yet been reported. We therefore performed IP of miniTurbo-Myc-MpSYP12A or miniTurbo-Myc-MpSYP13B using Myc-Trap beads from the same selected transgenic lines. After IP, we checked successful pulldown of the miniTurbo- Myc fusion-proteins by immunoblotting. We detected miniTurbo-Myc-MpSYP12A and miniTurbo-Myc-MpSYP13B fusion-proteins in cell extract that was used as input for IP, and in IP-eluates of samples from the transgenic lines (Fig. S3d). We then identified and quantified the proteins captured by Myc-IP using MS. We found a significant enrichment of the two bait proteins (Fig. 2a), confirming the immunoblotting result. Using Co-IP, we identified 4 and 1 potential interactors of MpSYP12A and MpSYP13B, respectively (Fig. 2a).

**Fig. 2.**
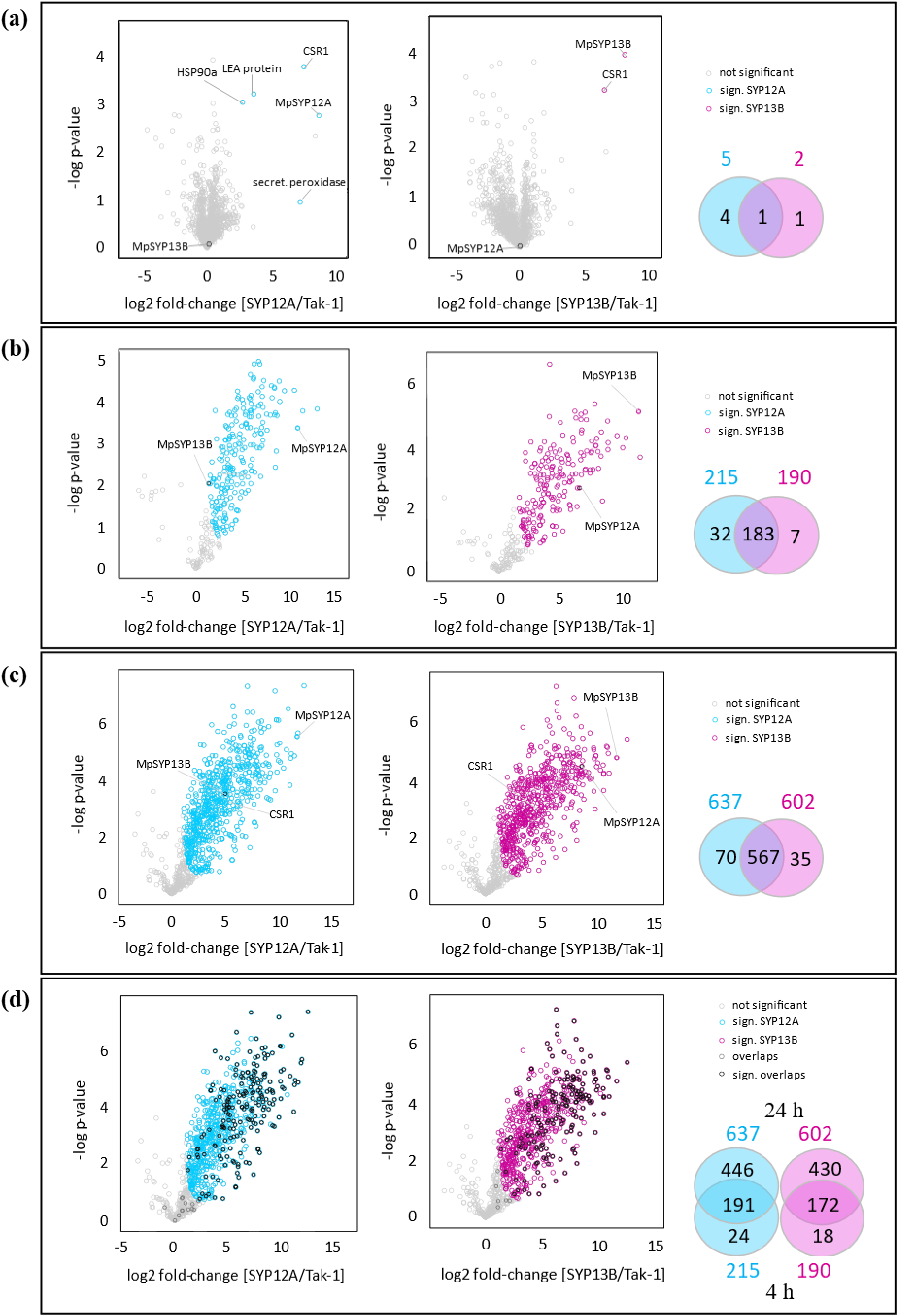
Identification of MpSYP12A or MpSYP13B interacting proteins by Co-IP and PL approaches. Wildtype Tak-1 was used as a control, and proteins that are significantly co-purified with or biotinylated by baits are highlighted. Potential interacting proteins for MpSYP12A and MpSYP13B are shown in turquoise and magenta, respectively. Venn diagrams show numbers of the potential interactors and their overlaps. (**a**) Myc-Trap Co-IP. Black text labels in volcano plots indicate the potential interactors for MpSYP12A or MpSYP13B. (**b**) 4 hours PL. (**c**) 24 hours PL. (**d**) Overlaps between 4 hours and 24 hours PL. Black circles in volcano plots indicate the potential interactors that are also identified with 4 hours PL. Dark grey circles indicate proteins that are identified as the potential interactors with 4 hours PL but not with 24 hours PL.

For the PL approach, we treated plants with biotin for 4 and 24 hours to identify and quantify proteins after pulldown by MS. As in the case of the Co-IP approach, we found a significant enrichment of both baits using the miniTurbo-based approach (Fig. 2b, c). Four hours of biotin treatment resulted in an identification of 214 and 189 proteins as potential interactors of MpSYP12A and MpSYP13B, respectively (Fig. 2b). By increasing the treatment time to 24 hours, the numbers of identified potential interactors nearly tripled (Fig. 2c). As expected, most of the candidates identified from 4 hours’ samples were also identified from 24 hours’ samples (Fig. 2d). In other words, approximately one third of the candidates that were identified after 24 hours of biotin treatment could be identified after 4 hours of treatment. MpCSR1 (CHLORSULFURON RESISTANT 1), a potential interactor of MpSYP12A and MpSYP13B that we identified using Co-IP, was identified in the 24 hours’ PL interactome dataset as well (Fig. 2c). Overall, these results demonstrate that PL approaches have a higher potential to identify undescribed interacting proteins compared to Co-IP approaches. However, it should be noted that the miniTurbo-based approach failed to identify proteins like secretory peroxidases that are predicted to be secreted into the extracellular space (Fig. 2a). This is reasonable as miniTurbo was fused to the intracellular domain of MpSYPs.

### PL using MpSYP12A and MpSYP13B as baits enriches proteins involved in vesicle-mediated transport and plasma membrane-localized proteins

We next asked whether the potential interactors that we identified by PL overall fit to the expected biological function of MpSYP12A and MpSYP13B. For this, we annotated *M. polymorpha* proteins based on information of *A. thaliana* homologs (Data S1). We then performed gene ontology (GO)- term enrichment analysis of the 24 hours interactome dataset using Metascape. Strikingly, ‘vesicle-mediated transport’ was the most significantly enriched GO-term extracted from the interactome data for both, MpSYP12A and MpSYP13B, coinciding with SNARE protein functions (Fig. 3a, b; Data S4). We also performed GO-term enrichment analysis of proteome data that were obtained by measuring the input samples used for the streptavidin-pulldown. Enriched GO-terms from proteome data were mainly related to primary metabolism (Fig. 3c), which is clearly distinct from the enriched GO-terms of the interactome dataset (Fig. 3a, b). These results suggest that the potential interactors comprise actual interactors of MpSYP12A and MpSYP13B. Analysis of the 4 hours interactome dataset gave similar results (Fig. S5a, b; Data S4).

**Fig. 3.**
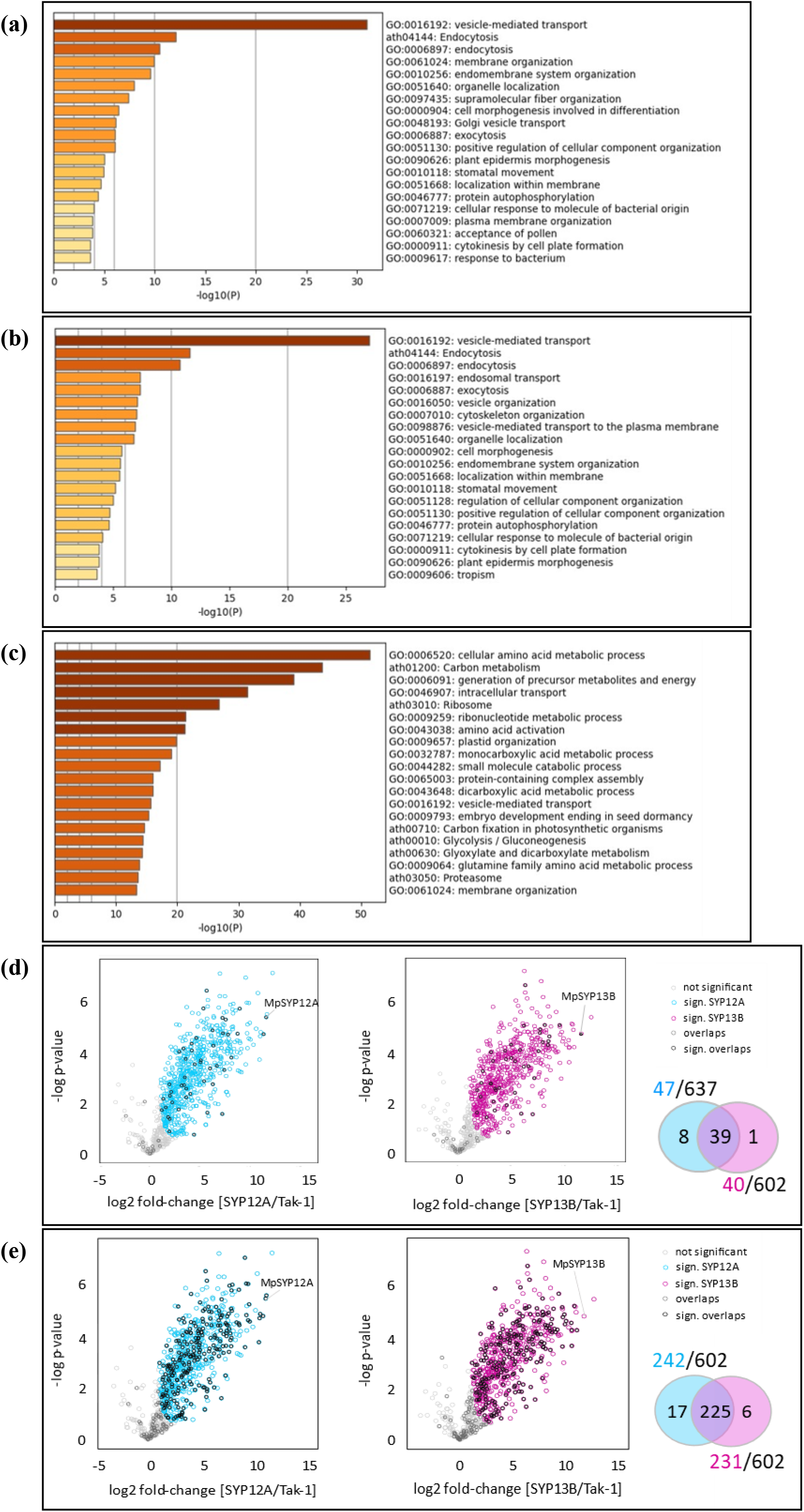
Features of the identified MpSYP12A or MpSYP13B interacting proteins. (**a** - **c**) GO-term enrichment analysis of (a) 24 hours PL MpSYP12A interactome, (b) 24 hours PL MpSYP13B interactome, and (c) measured whole proteome. The top 20 overrepresented GO-terms are shown. (**d**) *M. polymorpha* homologs of AtSYP1-interacting proteins are highlighted in black or grey on volcano plots of 24 hours PL. Venn diagram shows numbers of potential interactors that are homologous to the AtSYP1- interacors and their overlaps. (**e**) Potential interactors that are predicted to localize to the plasma membrane are highlighted in black or grey on volcano plots of 24 hours PL. Venn diagram shows numbers of the predicted plasma membrane-localized proteins and their overlaps.

A number of interactors of *A. thaliana* SNAREs, which are homologous to MpSYP12A and MpSYP13B, have been reported (Kwon *et al*., 2008; Fujiwara *et al*., 2014). Based on BioGrid, STRING, and IntAct databases, a total of 334 proteins were reported or predicted to interact with SYP1 proteins in *A. thaliana* (Data S2). Of these 334 *A. thaliana* proteins, we could identify 250 homologous proteins conserved in the *M. polymorpha* proteome using a BLAST approach (Data S3). Among these 250 proteins, we found that 47 and 40 proteins were identified as potential interactors of MpSYP12A and MpSYP13B, while 39 were shared between both baits, respectively. This means that around 11−15 % of all potential interactors of MpSYP12A and MpSYP13B revealed by the miniTurbo-based approach are homologous to known interactors of *A. thaliana* SYP1 proteins (Fig. 3d).

MpSYP12A and MpSYP13B were demonstrated to localize to the plasma membrane and to be ubiquitously expressed throughout the thallus in *M. polymorpha* (Kanazawa *et al*., 2016). By using PANTHER GO annotations and *A. thaliana* plasma membrane proteome data, we predicted plasma membrane localizations of the potential interactors in *M. polymorpha* (Data S1). We found that more than one third of the potential interactors are expected to be plasma membrane- localized (Fig. 3e). This result further supported the specificity and utility of the miniTurbo-based approach.

### The miniTurbo-mediated approach can reveal subtle differences between very similar baits

By comparing the 24 hours interactome data for MpSYP12A and MpSYP13B, we found that both baits share 90−95 % of their potential interactors (Fig. 1c), which is reasonable considering their predicted functions. Still, we captured 52 and 9 proteins that preferentially interact with MpSYP12A and MpSYP13B, respectively (Fig. 4a, Table 1). To investigate exclusive interaction partners of MpSYP12A and MpSYP13B, we implemented iBAQ (Intensity Based Absolute Quantification) values to the volcano plot (Fig. 4b). The iBAQ is a measure of protein abundance, and relative abundance was reflected in different circle sizes. The respective colours indicate abundances in all pulldown samples. By this, we revealed that MpNEK potentially interacts exclusively with MpSYP13B, and we found 13 proteins that potentially interact exclusively with MpSYP12A (Fig. 4b, Table 1).

**Fig. 4.**
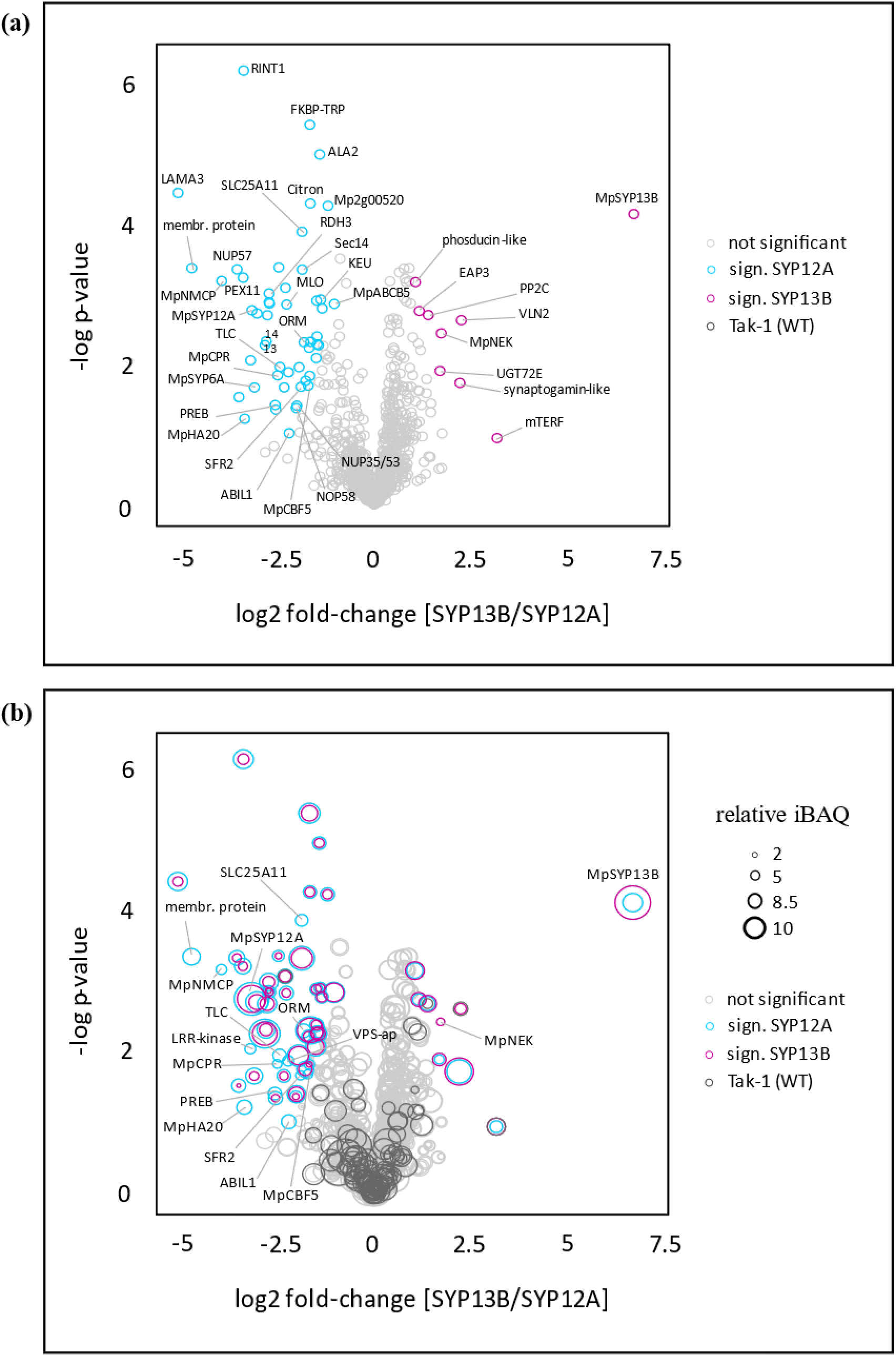
Potential interactors that preferentially interact with MpSYP12A or MpSYP13B. (**a**) Proteins that preferentially interacted with MpSYP12A or MpSYP13B are highlighted in turquoise and magenta, respectively. (**b**) Relative protein abundances based on iBAQ values are indicated by sizes of circles. Protein abundances in the samples of MpSYP12A, MpSYP13B, and wildtype Tak-1 are shown in turquoise, magenta, and dark grey, respectively. Proteins that were exclusively identified from the samples of MpSYP12A or MpSYP13B but not from the wildtype Tak-1 sample are annotated except for MpSYP12A and MpSYP13B.

**Table 1.**
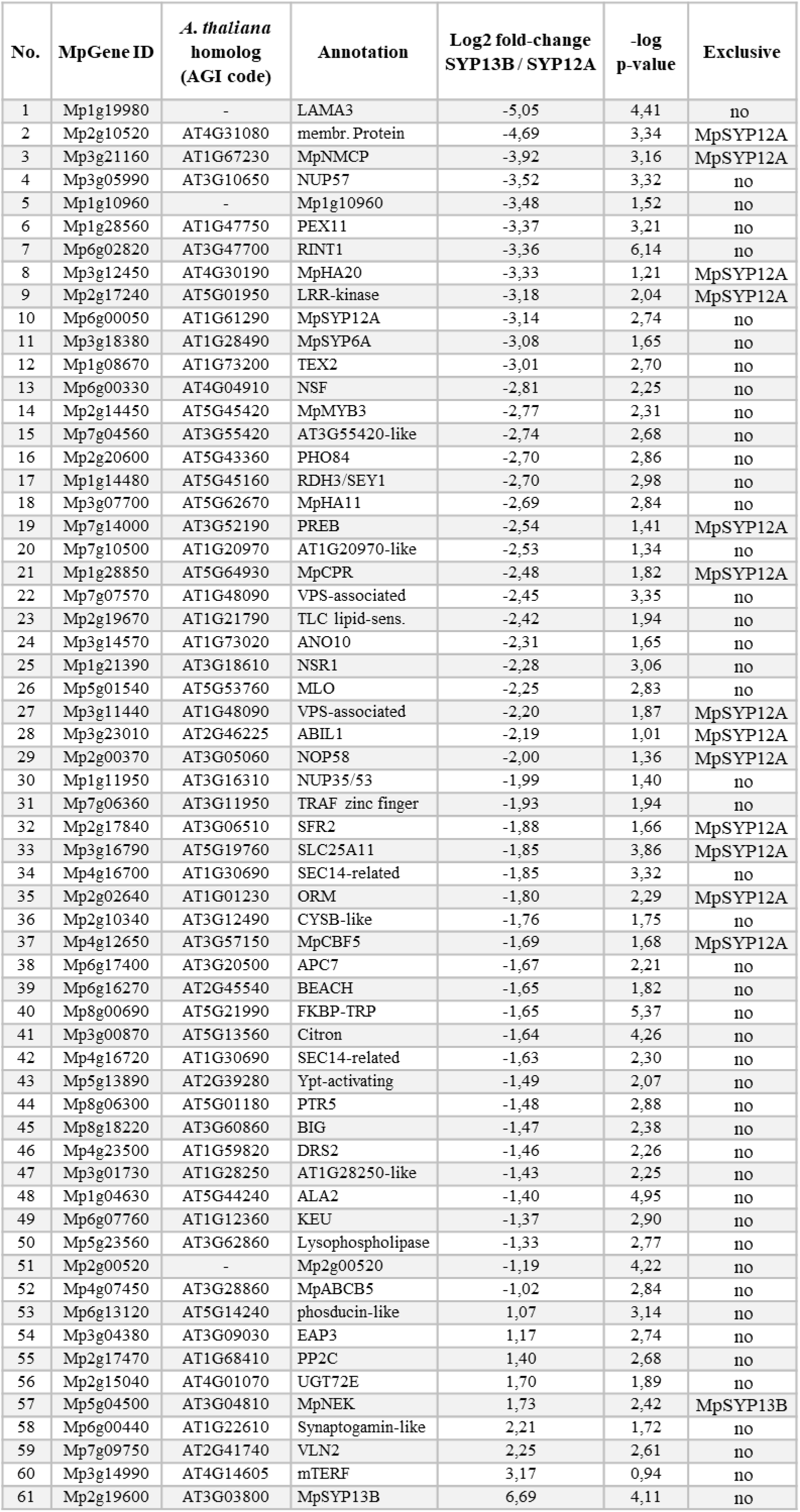
Potential interactors that preferentially interact with MpSYP12A or MpSYP13B. Proteins that are highlighted in turquoise or magenta in Fig. 4 are listed.

## Discussion

Published studies applying *in planta* TurboID-based PL have utilized model angiosperm species, and the amenability of current PL approaches to other plant species remains to be determined as outlined by Mair and Bergmann (2021). Here, we successfully applied miniTurbo-based PL in the liverwort *M. polymorpha*. This confirmed the transferability of biotin ligase utilized interactomics to a model bryophyte species.

We tested different biotin concentrations and treatment times to investigate biotin-labelling of proteins in *M. polymorpha* cells. Mair *et al*. (2019) reported a saturation of biotinylated proteins in *A. thaliana* stable transgenic lines after treatment with 50 µM biotin solution for 1 hour. Zhang *et al*. (2019) transiently expressed TurboID-fusion-proteins in *N. benthamiana* leaves and tested biotin concentrations up to 800 µM. Zhang *et al*. observed that protein biotinylation was saturated after 8 hours of treatment with 200 µM biotin, and that 15 minutes were sufficient for the saturation at 200 µM biotin concentration. Meanwhile, in *M. polymorpha*, we did not observe a saturation of protein biotinylation after 24 hours of treatment with 700 µM biotin. The observed differences in saturation of biotinylated proteins among the different studies could be partially explained by differences in mechanisms and efficiencies for biotin uptake and metabolism. Other factors potentially impacting biotin-labelling activities are plant growth conditions like light cycle and temperature used during biotin treatment.

For PL approaches, a careful evaluation of false positive candidates based on well-designed controls is desirable to generate a set of candidates with high confidence for further analyses and validation. It would be beneficial to include other plasma membrane-localized proteins that are independent of MpSYP12A and MpSYP13B to aid in predicting false positive candidates, which might have been biotinylated randomly due to their localizations. That is to say, our transgenic lines expressing miniTurbo-Myc-MpSYP12A and miniTurbo-Myc- MpSYP13B could be used as suitable controls for interactome mapping using other plasma membrane-localized baits in the future. Meanwhile, the overall high similarity between MpSYP12A and MpSYP13B can be exploited to investigate specific interactors to understand functional differences between the two SNAREs. In general, long biotin treatment times potentially increase false positive labelling. On the other hand, longer treatment times may be required to efficiently capture rare or transient interactors.

For biotin-labelling approaches, protein extraction can be conducted in the presence of strong detergents. Strong detergents facilitate the extraction and solubilization of membrane-associated proteins, which can be advantageous for interactomics of plasma membrane-localized or organellar proteins. Our results confirmed that we could indeed capture a manifold of proteins that are predicted to be plasma membrane- localized, which has not been tested on the proteome level in published studies using PL in plants. In contrast to the Co-IP approach, the miniTurbo-based method could not capture extracellular interactors of MpSYP12A and MpSYP13B, which can be a drawback for certain applications. In the future, it remains to be determined whether the miniTurbo-based approach is also suitable for other cellular compartments, which may have a different intraorganellar pH or temperature, as discussed by Mair and Bergmann (2021).

Until to date, a manifold of interaction partners of AtSYP1 family proteins has been identified or predicted (Kwon *et al*., 2008; *Arabidopsis* Interactome Mapping Consortium, 2011; Fujiwara *et al*., 2014);. With our miniTurbo-based approach, we were able to capture homologs to well-known AtSYP1-interacting proteins, such as KEU, NPSN11, SYP61, and VAMP721 (Fujiwara *et al*., 2014), demonstrating the reliability of our method. We identified 47 and 40 proteins that are homologous to previously described AtSYP1-interacting proteins, while 104 and 92 proteins were linked to ‘vesicle- mediated transport’ based on our GO-term enrichment analysis. This indicates that we were able to capture a number of undescribed potential interactors of MpSYP12A and MpSYP13B, respectively. Kanazawa *et al*. (2020) were able to show that MpSYP12A but not MpSYP13B localized to the phragmoplast during cell plate formation by using fluorescent reporter-lines. Based on our GO-term enrichment analysis, we found the same set of 30 proteins related to ‘cell plate formation’ in interactomes of both, MpSYP12A and MpSYP13B. In other words, we failed to identify unique potential interactors of MpSYP12A previously linked to ‘cell plate formation’. To address a role of MpSYP12A at the phragmoplast, experimental conditions would need to be further adjusted to capture the interactome during cell division, by using cell cycle specific promoters to express proteins in dividing cells, for instance.

Our miniTurbo-based approach identified a number of potential interactors that are specific for MpSYP12A and MpSYP13B, which may help to understand functional differences between the two SNAREs in the future. For example, we found that homologs to MILDEW RESISTANCE LOCUS O (MLO) and PENETRATION 3 (PEN3) preferentially interact with MpSYP12A. *MLO* genes were shown to be involved in susceptibility to powdery mildew pathogens in barley (Büschges *et al*., 1997) and *A. thaliana* (Consonni *et al*., 2006). At*PEN3* was found to be involved in resistance against barley powdery mildew (Stein *et al*., 2006) and cell-death responses upon infection with *P. infestans* (Kobae *et al*., 2006). In *A. thaliana*, AtSYP121 or PENETRATION1 (PEN1) is homologous to MpSYP12A.

At*PEN1* was shown to be required for SNARE-dependent penetration resistance against barley powdery mildew and pathogen-induced vesicle accumulation was enhanced in *MLO* loss-of-function mutants (Collins *et al*., 2003). Recently, Rubiato *et al*. (Rubiato *et al*., 2021) provided evidence for an evolutionary conserved role of SYP12 proteins in the formation of papillae and encasements at pathogen penetration sites, which are effective defence structures against a broad range of filamentous pathogens. Taken together, our data suggest a function of MpSYP12A in penetration resistance and responses to filamentous pathogens, like SYP12 proteins in other plant species. Besides, we found that MpSYP12A may exclusively interact with ABSCISIC ACID INSENSITIVE1 (MpABI1), NUCLEAR MATRIX CONSTITUENT PROTEIN (MpNMCP), and homologs of OROSOMUCOID-LIKE 1 (ORM1) and SENSITIVE TO FREEZING 2 (SFR2). MpABI1 is involved in abscisic acid signalling (Tougane *et al*., 2010) and MpNMCP was found to function in stress signalling in *M. polymorpha* (Wang *et al*., 2021). In *A. thaliana*, NMCP homologs play a role in immunity (Choi *et al*., 2019; Jarad *et al*., 2019). AtORMs were reported to play roles in sphingolipid homeostasis and stress responses (Li *et al*., 2016), and AtSFR2 is a membrane remodelling enzyme responsive to freezing conditions in *A. thaliana* (Barnes *et al*., 2019). Accordingly, our findings may suggest a potential role of MpSYP12A in lipid homeostasis and stress responses. We identified NIMA- related protein kinase 1 (MpNEK) as potential exclusive interactor for MpSYP13B. MpNEK directs tip growth in rhizoids of *M. polymorpha* (Otani *et al*., 2018), and thus our data may suggest a potential role of MpSYP13B in rhizoid tip growth. It should be noted that potential interactors revealed by PL need to be validated using complementary approaches. In summary, our interactome data should be a useful resource for future investigations of functional conservation and diversification of SNARE proteins in plants.

## Acknowledgements

We thank Takashi Ueda (National Institutes of Natural Sciences, Division of Cellular Dynamics, Tokyo, Japan) for providing the plasmids harbouring genomic sequences for native expression and regulation of MpSYP12A and MpSYP13B. We thank Takayuki Kohchi (Kyoto University, Japan) for providing pMpGWB vectors.

## Author Contribution

KM and HN have designed the research. KM, SCS, AH, and HN have contributed to experimental design and workflow conceptualization. KM, SCS, and AH have conducted experiments. KM, SCS, and HN have analysed the data by mass spectrometry. KM, SCS, and HN wrote the manuscript. This project was supported by the Max Planck Society and was conducted in the framework of MAdLand (http://madland.science, Deutsche Forschungsgemeinschaft (DFG) priority program 2237). HN is grateful for funding by the DFG (NA 946/1-1).

## Data Availability

The mass spectrometry proteomics data have been deposited to the ProteomeXchange Consortium via the PRIDE [1] partner repository with the dataset identifier PXD030429.

---

**Fig. S1.**
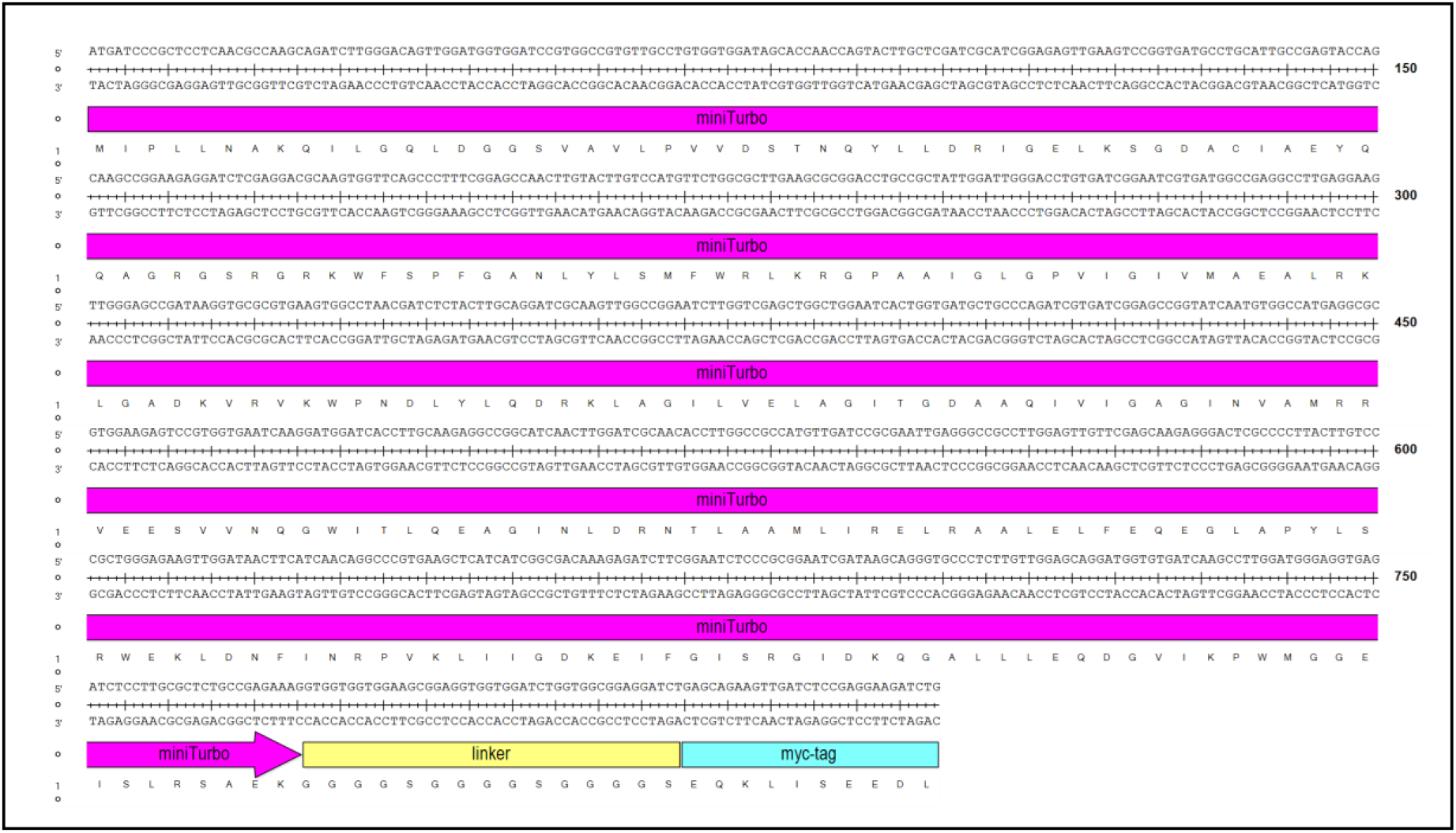
Nucleotide and amino acid sequence of the miniTurbo biotin ligase used in this study. The nucleotide sequence of the original miniTurbo (Branon et al., 2018) was modified based on preferential codon-usage of *M. polymorpha*. A linker sequence of 15 amino acids length was added to the C-terminus of miniTurbo, to minimize potential sterical impairments of the enzymatic function. The linker sequence was fused to a single Myc-tag peptide of 10 amino acids length, to enable immunoblot detection of miniTurbo-fusion proteins and affinity purification using Myc-Trap.

**Fig. S2.**
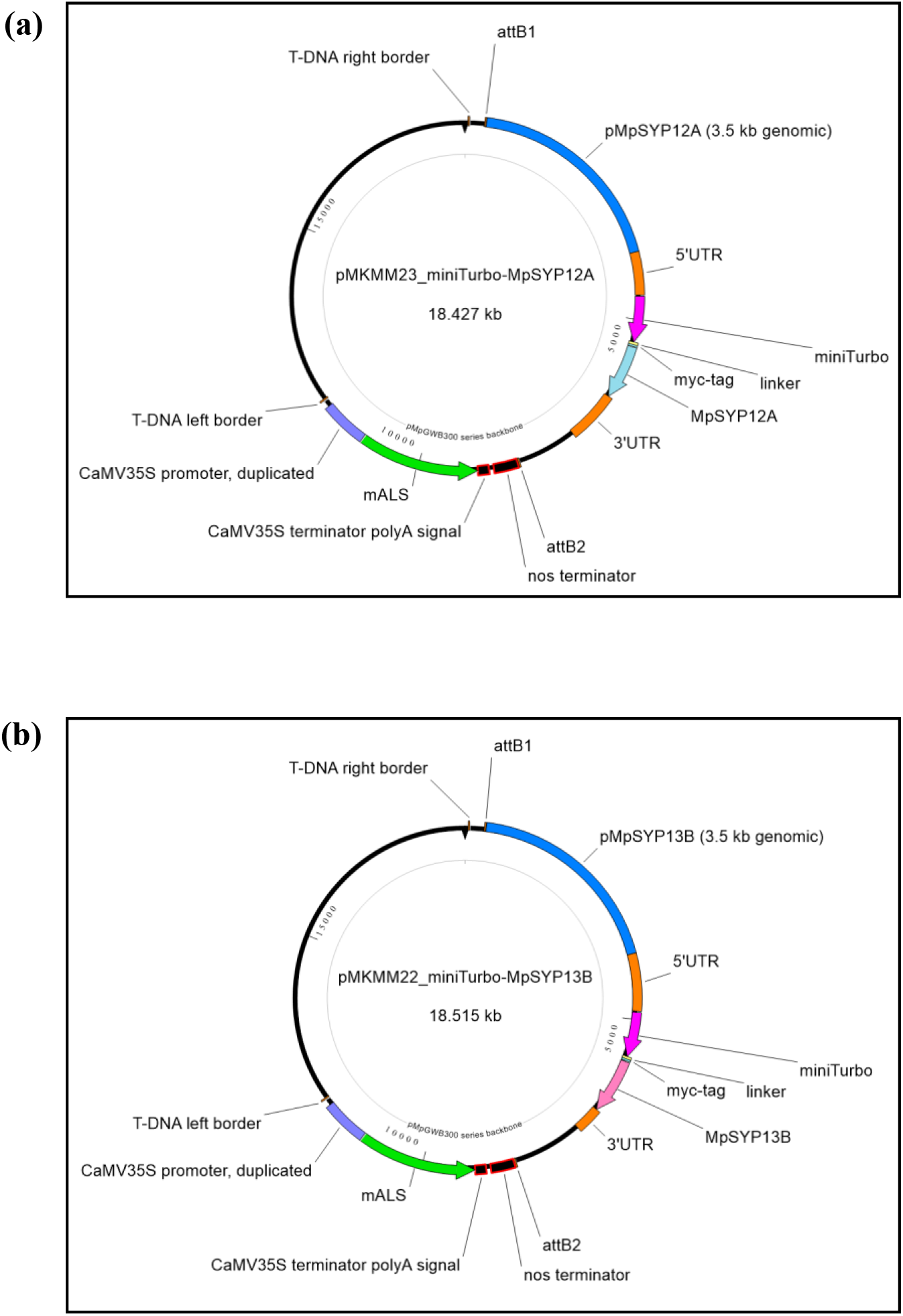
Plasmid maps of the binary vectors used in this study. (**a**) Vector map of pMKMM23 for the expression of miniTurbo-Myc-MpSYP12A. (**b**) Vector map of pMKMM22 for the expression of miniTurbo-Myc- MpSYP13B.

**Fig. S3.**
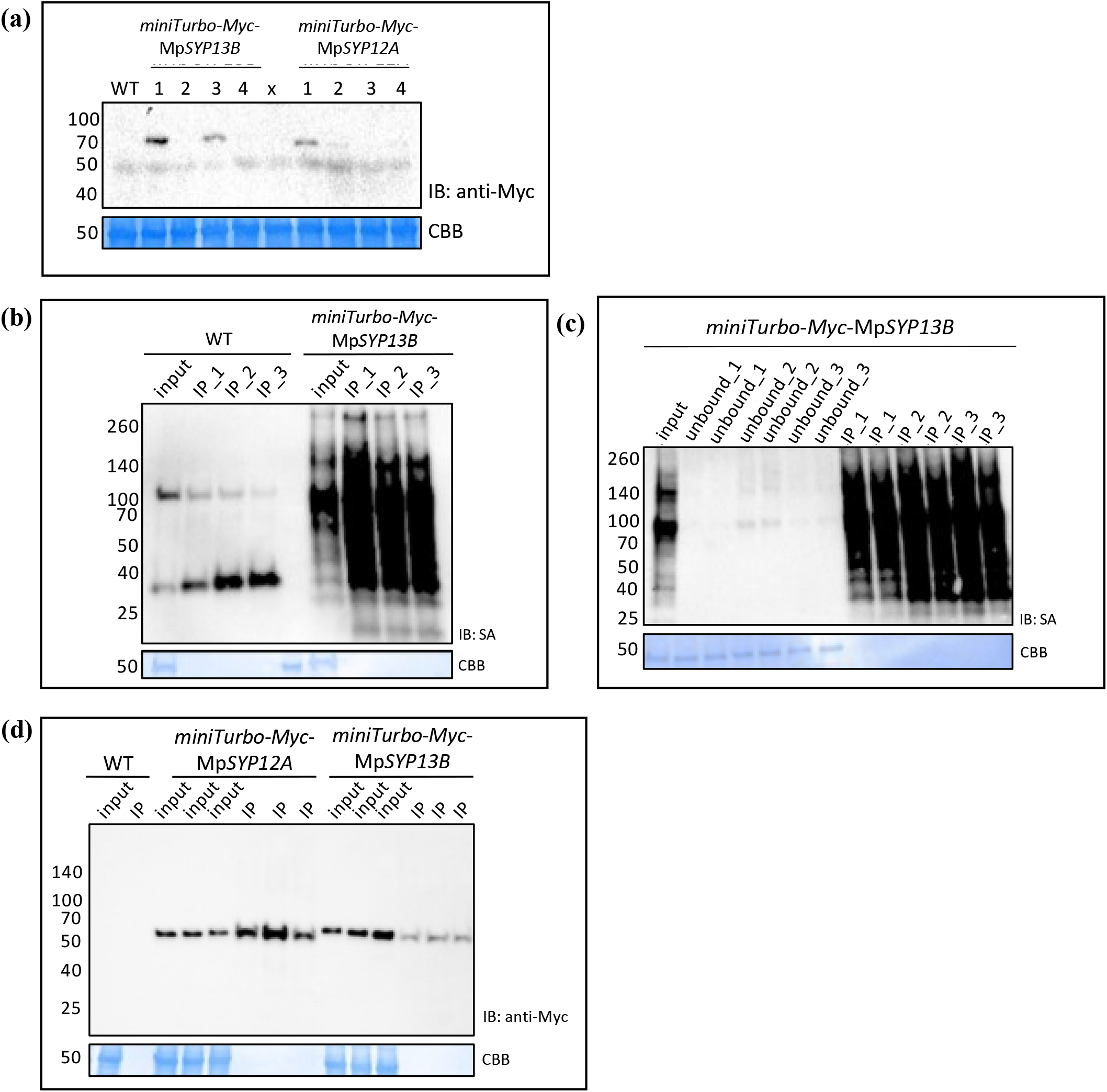
Evaluation of the fusion protein expression, biotin depletion methods, and Myc-Trap Co-IP by immunoblotting. (**a**) Selection of transgenic lines expressing miniTurbo-Myc-MpSYP12A and miniTurbo- Myc-MpSYP13B fusion-proteins. Immunoblot (IB) of cell extracts from 10- day old thalli. MiniTurbo-Myc fusion proteins were detected by using an anti-Myc antibody. Cell extract of wildtype Tak-1 (WT) was used as a control. (**b** and **c**) Comparison of biotin depletion methods. Ten-day old thalli of a transgenic line expressing miniTurbo-Myc-MpSYP13B and wildtype Tak-1 (WT) were treated with 700 µM biotin solution for 24 hours. Total protein was extracted (input) and free biotin was removed from the samples by methanol:choloform precipitation (IP_1), PD-10 column desalting (IP_2), or Zeba spin column desalting (IP_3) before pulldown of biotinylated proteins using streptavidin-agarose beads. Streptavidin (SA) Immunoblots (IB) of biotinylated proteins. (**b**) Biotinylated proteins in cell extracts (input) and IP-eluates (IP-1, IP_2, IP_3) for the different biotin depletion methods. (**c**) Biotinylated proteins in cell extract (input), the supernatant of the streptavidin-agarose beads after affinity pulldown (unbound_1, unbound_2, unbound_3), and IP-eluates (IP_1, IP_2, IP_3) for the different biotin depletion methods. All depletion methods were tested in duplicates. (**d**) Evaluation of Myc-Trap Co-IP by immunoblotting. Immunoblot (IB) of cell extracts from 10-day old thalli. MiniTurbo-Myc fusion proteins were detected by using an anti-Myc antibody in cell extracts (input) and IP-eluates (IP) after affinity-purification using Myc-Trap beads. Coomassie Brilliant Blue-stained (CBB) membranes are shown as loading controls.

**Fig. S4.**
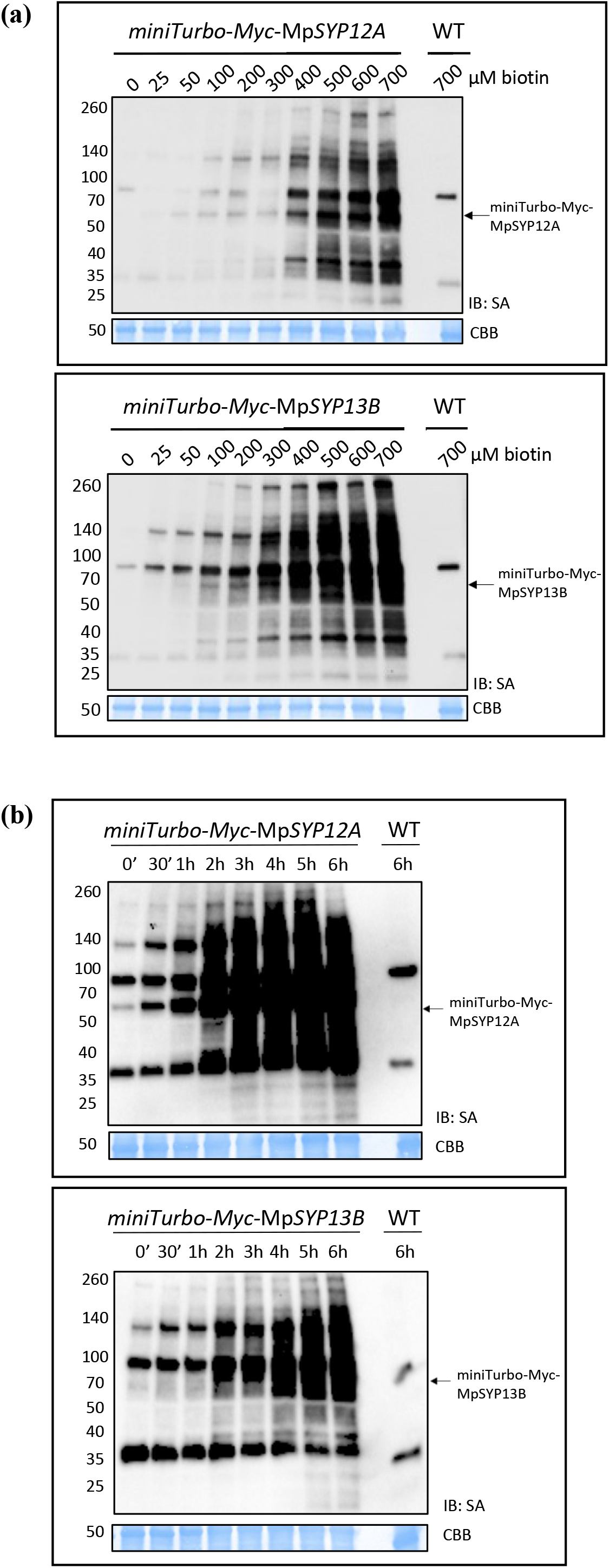
Biotin ligase activity in *M. polymorpha*. Streptavidin (SA) immunoblots (IB) of cell extracts from *M. polymorpha* transgenic lines expressing miniTurbo-Myc-MpSYP12A (upper panels) and miniTurbo- Myc-MpSYP13B (lower panels) that were (**a**) treated with 0−700 µM biotin solutions for 24 hours at RT or (**b**) treated with 700 µM biotin solution for 0−6 hours at RT. Cell extracts of wildtype Tak-1 (WT) treated with 700 µM biotin solution for (**a**) 24 hours or (**b**) 6 hours were used as controls. Arrows indicate the positions of the biotinylated miniTurbo-Myc-MpSYP12A and miniTurbo-Myc-MpSYP13B fusion-proteins. Coomassie Brilliant Blue- stained (CBB) membranes are shown as loading controls.

**Fig. S5.**
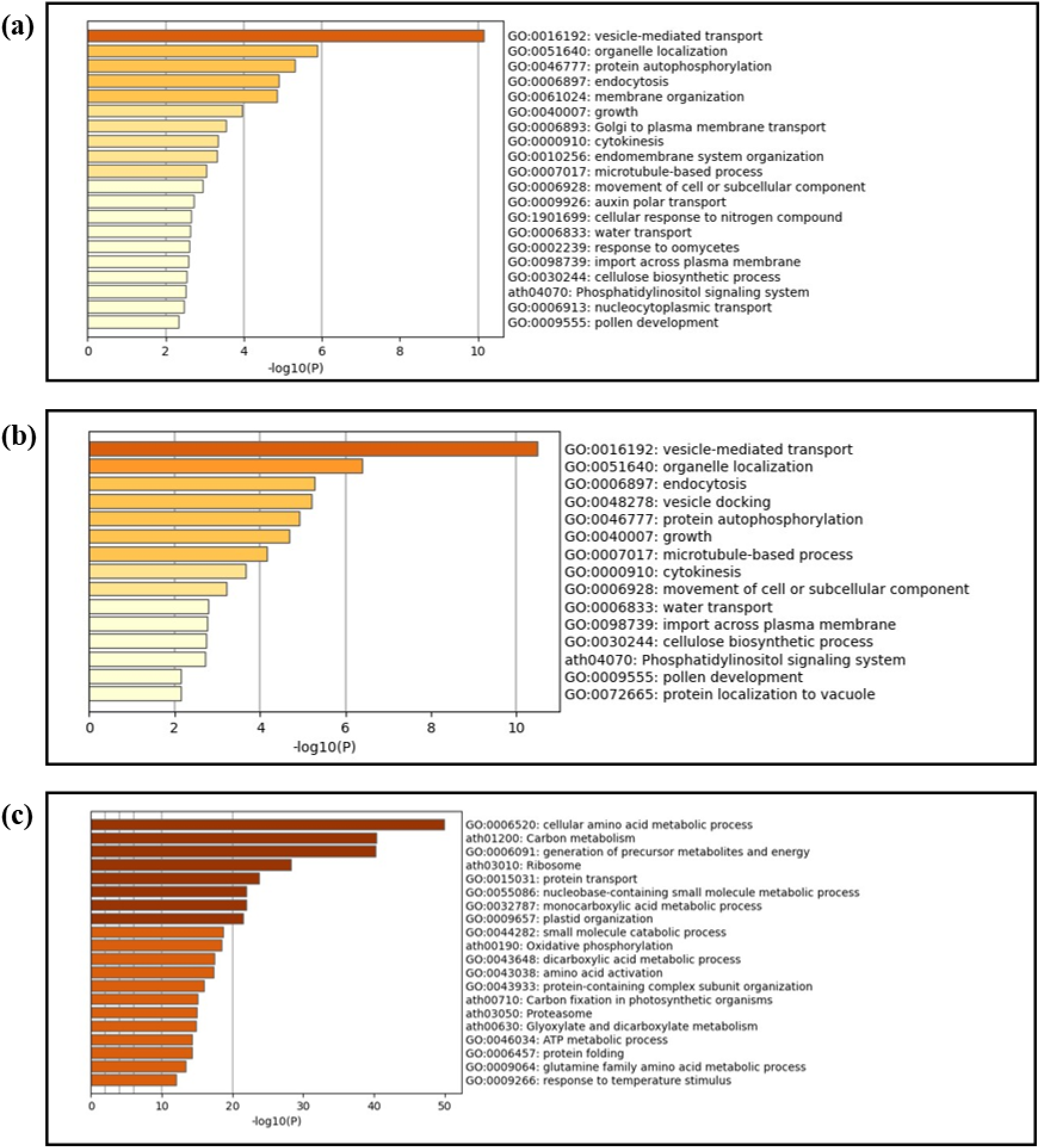
GO-term enrichment analysis. GO-term enrichment analysis of (a) 4 hours PL MpSYP12A interactome, (b) 4 hours PL MpSYP13B interactome, and (c) measured whole proteome. The top 20 overrepresented GO-terms are shown.

**Fig. S6.**
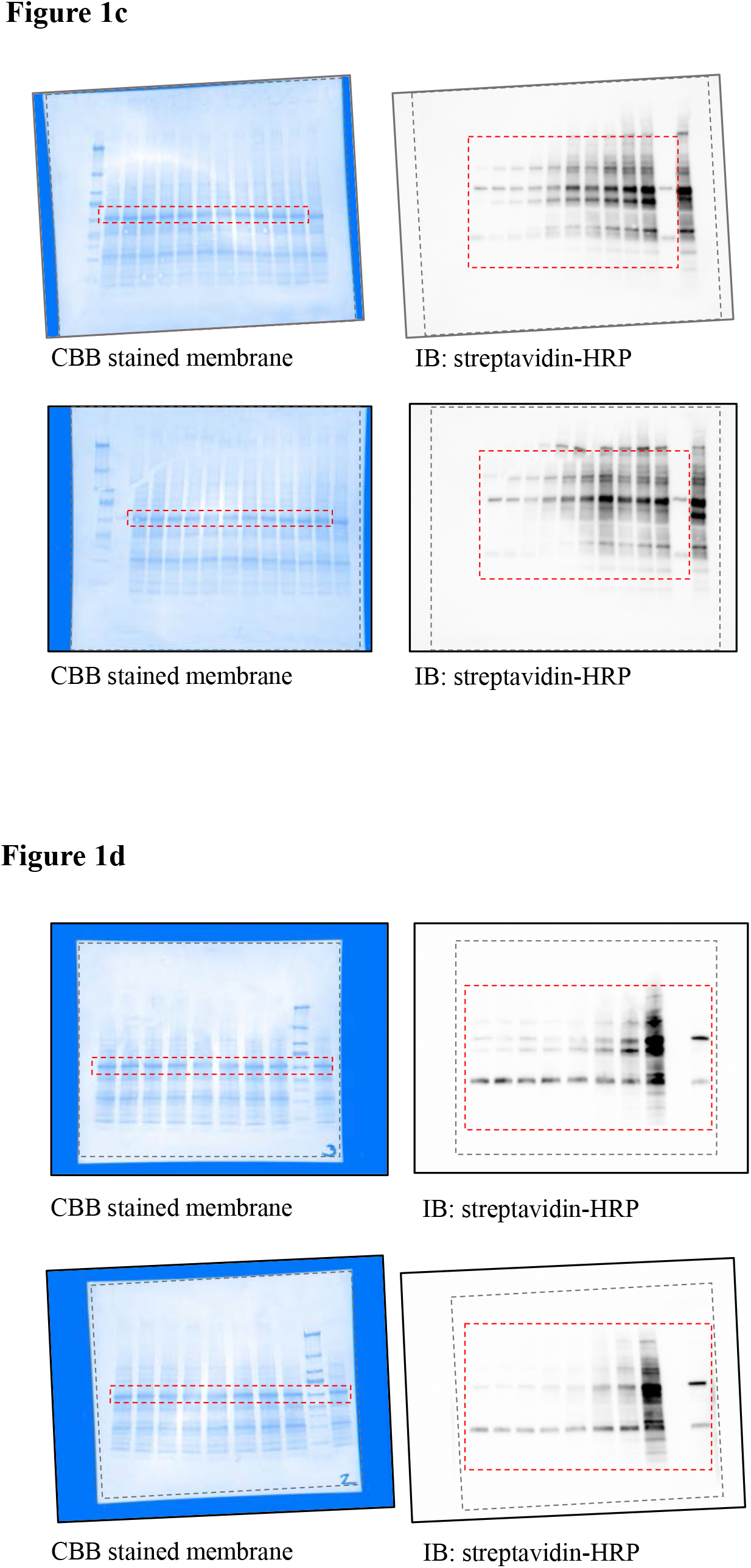
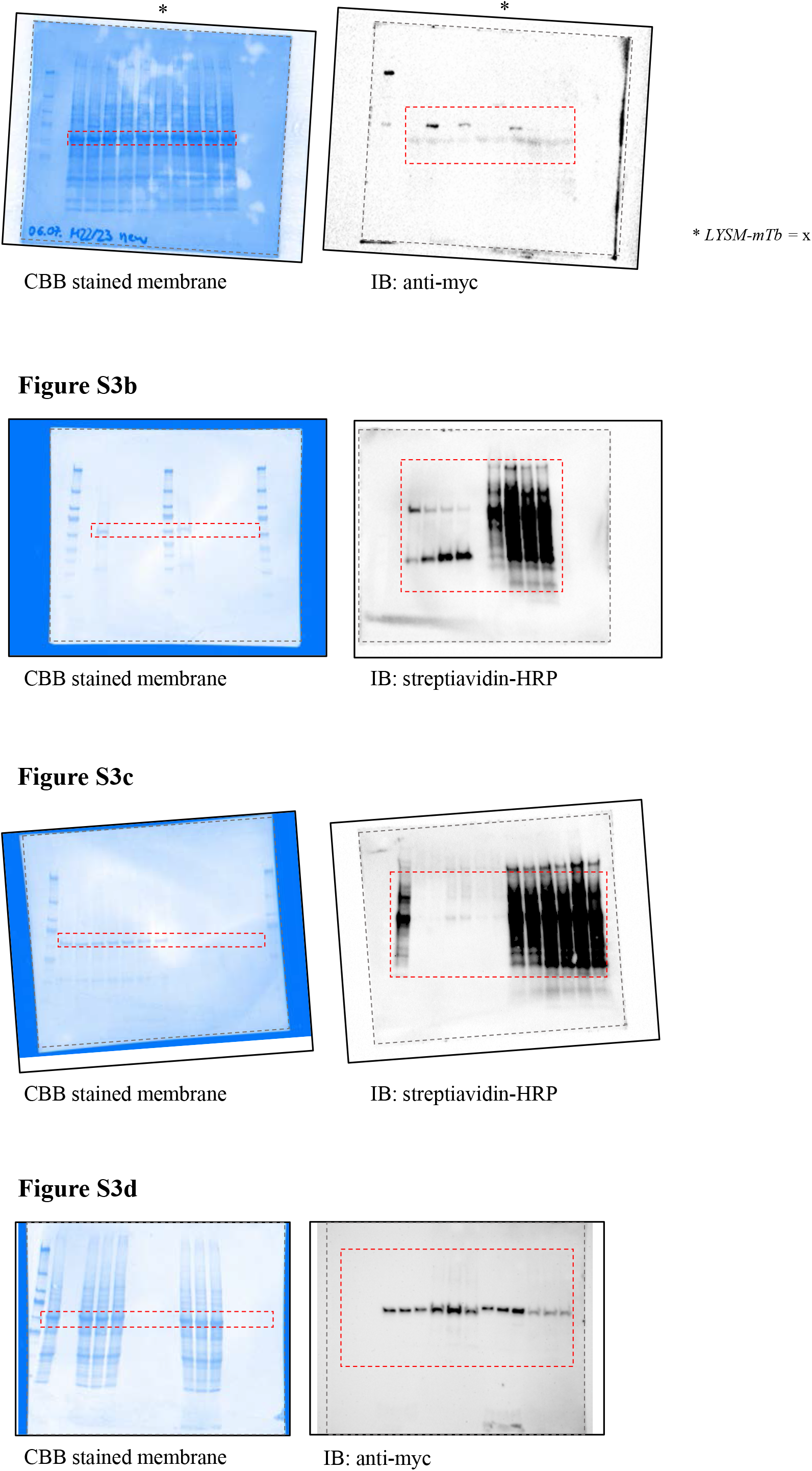
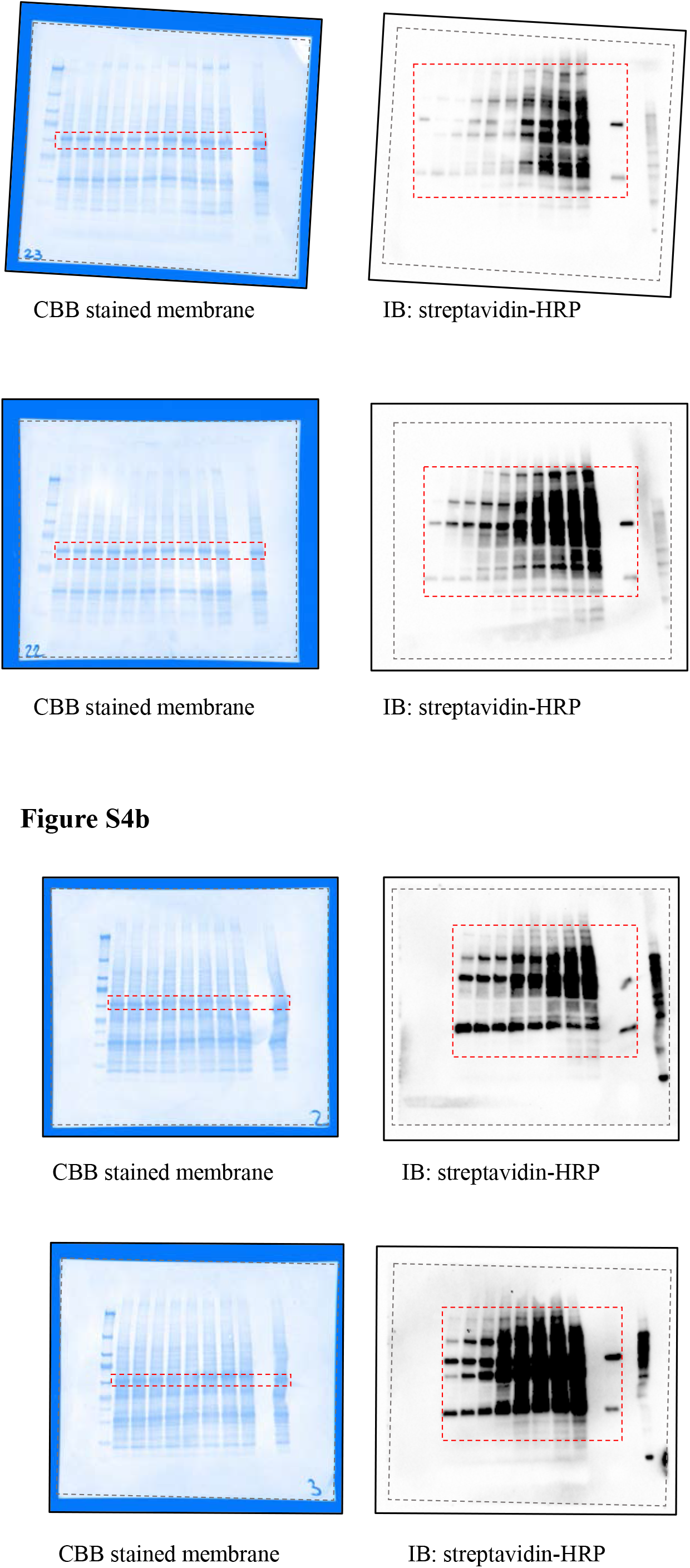
Uncropped images of immunoblots used in figures.

